# The Cuon Enigma: Genome survey and comparative genomics of the endangered Dhole (*Cuon alpinus*)

**DOI:** 10.1101/443119

**Authors:** Bilal Habib, Pallavi Ghaskadbi, Parag Nigam, Shrushti Modi, Peddamma Sathish Kumar, Kanika Sharma, Vidhya Singh, Bipin Kumar, Abhishek Tripathi, Harish Kothandaraman, Sailu Yellaboina, Dushyant Singh Baghel, Samrat Mondol

## Abstract

The Asiatic wild dog is an endangered monophyletic canid restricted to Asia; facing threats from habitat fragmentation and other anthropogenic factors. Dholes have unique adaptations as compared to other wolf-like canids for large litter size (larger number of mammae) and hypercarnivory making it evolutionarily notable. Over evolutionary time, dhole and the subsequent divergent wild canids have lost coat patterns found in African wild dog. Here we report the first high coverage genome survey of Asiatic wild dog and mapped it with African wild dog, dingo and domestic dog to assess the structural variants. We generated a total of 124.8 Gb data from 416140921 raw read pairs and retained 398659457 reads with 52X coverage and mapped 99.16% of the clean reads to the three reference genomes. We identified ~13553269 SNV’s, ~2858184 InDels, ~41000 SVs, ~1854109 SSRs and about 1000 CNVs. We compared the annotated genome of dingo and domestic dog with dhole genome sequence to understand the role of genes responsible in pelage pattern, dentition and mammary glands. Positively selected genes for these phenotypes were looked for SNP variants and top ranked genes for coat pattern, dentition and mammary glands were found to play a role in signalling and developmental pathways. Mitochondrial genome assembly predicted 35 genes, 11 CDS and 24 tRNA. This genome information will help in understanding the divergence of two monophlyletic canids, *Cuon* and *Lycaon*, and the evolutionary adaptations of dholes with respect to other canids.

## Introduction

The Asiatic wild dog (*Cuon alpinus*) or dhole is monotypic canid belonging to the genus Cuon. Dholes were widely distributed across the continents of North America, Europe and Asia during the Pleistocene era (Cohen, 1978), but are now restricted to parts of South, East and Southeast Asia (Kamler et al. 2015). Threats like habitat loss, diseases from domestic dogs, persecutions and interspecific competition among large co-predators (Durbin et al. 2004) have resulted in a 75% decline of the historic range of dholes (Kamler et al. 2015). The remnant populations of dholes in Asia are highly fragmented (Burton 1940, Durbin et al 2004, Karanth and Sunquist 2000) with a decreasing population trend placing them under the ‘Endangered’ category of the IUCN (category C2a(i)) (Kamler et al. 2015) and ‘Appendix II’ of the Convention on International Trade in Endangered Species (CITES). The Indian subcontinent currently harbours majority of the wild dhole populations across its range (Kamler et al. 2015), where the species has experienced about 60% habitat decline (Karanth et al. 2010). The dense forests of Western Ghats and central India retain most of the dhole population (Karanth et al. 2009) while the Eastern Ghats landscape, northeast India and Himalayan region hold smaller populations (Karanth et al. 2009, Lyngdoh et al. 2014, Bashir et al. 2014).

Ecologically, dhole is the only social canid in closed forest systems across their range. In India, dhole shares habitat with larger co-predators like tigers and leopards (Karanth and Sunquist 2000). In spite of smaller body size, their group living and pack-hunting strategies help them survive intense intraguild competition (Wang and Macdonald 2009; Steinmetz et al. 2013). Dholes are popularly known as ‘whistling hunters’ due to the use of whistles as a unique mode of communication among large packs (Fox 1984). They also have other specialized communication mechanisms such as varied scent-marking behaviours which include cross-marking ritual by dominant males and females- a form of mate guarding, body-rubs and hand-stand scent marking (Ghaskadbi et al. 2016).

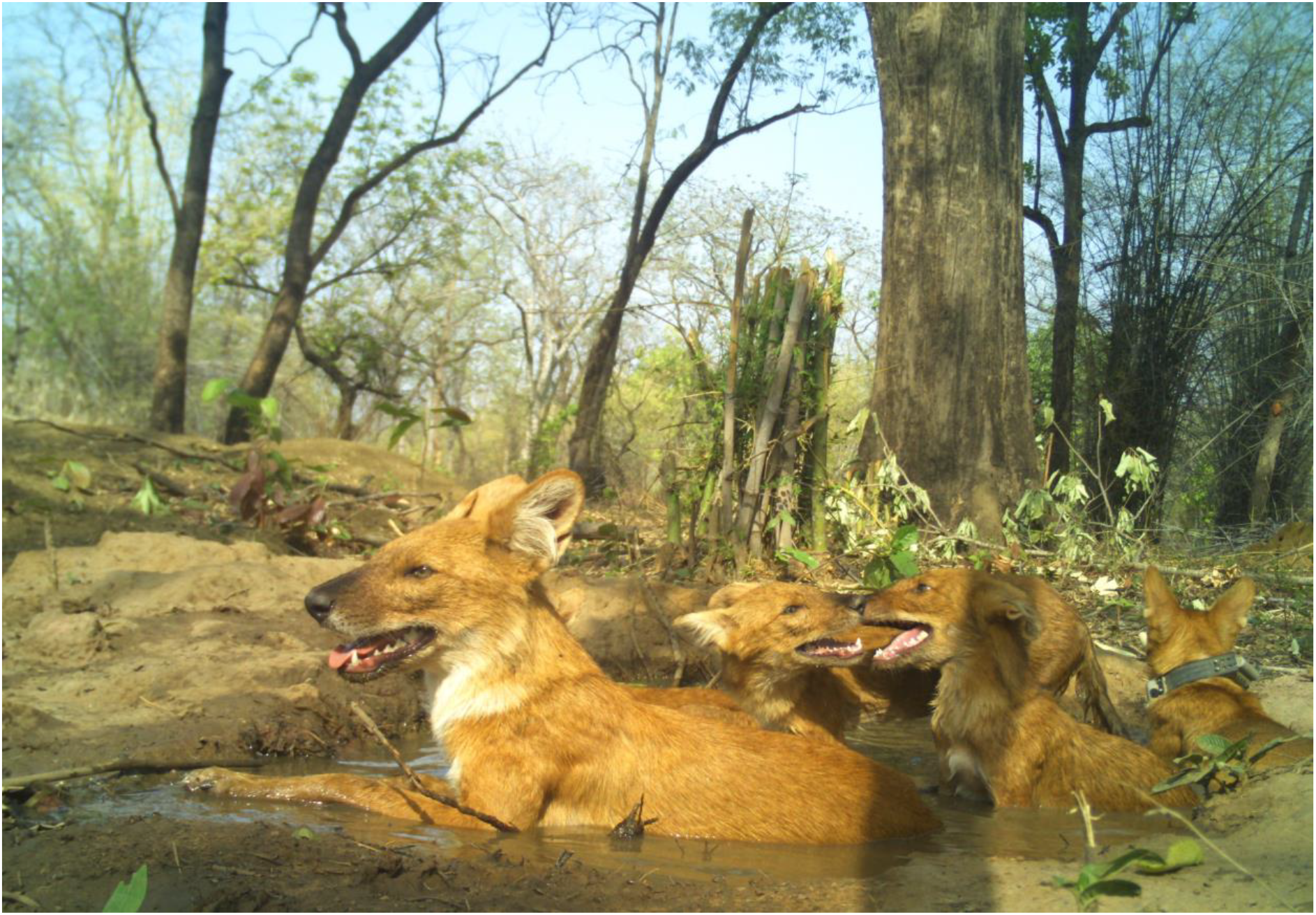

Evolutionarily, dholes are grouped with other wolf-like canids (Gray wolf, coyote and Ethiopian wolf) (Wayne et al. 1997) and diverged within this group approximately 4-6 million years ago after the African wild dog (Lindblad-Toh et al. 2005). Both species evolved with a distinctly structured unicuspid talonid on the lower carnassial for enhanced meat-shearing capacity (Van Valkenburgh, 1991) and 6-7 pairs of mammae, possibly for parental care of large litter size (Burton, 1940). However, dhole and the subsequent divergent wild species in this clade (Ethiopian wolf, coyote, gray wolf, golden jackal, black-backed jackal and side-striped jackal) have lost the characteristic pelage pattern that is only present in African wild dog. All the above species-specific unique adaptations of dholes make them an evolutionary enigma. Given the current anthropogenic pressures, the survival of this monotypic genus depends on an integrated approach of conservation measures involving detailed information on ecology, demography and genetics.

In this paper, we report the first genome survey of wild-ranging Asiatic wild dog, where we mapped it against the closely related African wild dog (*Lycaon pictus*), dingo (*Canis lupus dingo*) and domestic dog (*Canis familiaris*). Further, we describe dhole genome structural variants, copy number variants, and simple sequence repeats and conduct mitochondrial genome assembly and annotation. Finally, we also explore dhole-specific variations in coat colour, dentition and mammary gland gene complexes. This genomic data would help in future studies on dhole evolution and demographic history.

## Methods

### Sample collection, permits and ethical considerations

As part of an ongoing study in the Tadoba Andhari Tiger Reserve (Permit No. D-22(8)/WL/Research/CT-722/(12-13)/2934/2013:), five free-ranging dholes were captured from the wild and radio-collared for intensive monitoring (Permit No. SPP-12/2016 and SPP-22/2017). Out of the five individuals, the blood sample of one adult male was used for genome sequencing. The dholes were highly social and active making them difficult targets to capture in wild. We immobilized the target individuals using Zoletil (Zoletil 100; Virbac, Carros, France) (Sawas et al. 2005). The samples were stored in EDTA vacutainers at −20 0C for further analysis.

### Library construction, sequencing and filtering

Genomic DNA was extracted from the blood sample with Nucleospin Blood Kit (MACHAREY-NAGEL Gmbh & Co. KG, Duren, Germany). Lysis was performed with 200μl of lysis buffer (B3), 25μl of Proteinase K for 200μl of blood, followed by the manufacturer’s protocol provided in the kit. DNA was eluted with 100 μl of 1X TE, and stored at −20 ^0^C for further analysis.

Paired end libraries were constructed using NEBNext^®^ UltraTM DNA Library Prep Kit for Illumina^®^ following manufacturer’s protocols. These libraries were prepared with insert size of 300bp and 500bp using 1 μg of initial genomic DNA. The PCR products were purified to optimize the size of the fragments using AMPure XP system, and fragments were selected based on size using Agilent 2100 Bioanalyzer. Each insert library was run in multiple lanes using an Illumina HiSeq 2500 till optimal coverage for analysis was achieved. We generated a total of 124.8 Gb data from 416140921 raw read pairs, leading to approximately 52x coverage based on a genome size estimation of roughly 2.4 Gb. We trimmed the low-quality bases from both the sides using Trimmomatic v 0.36 (Bolger et al. 2014) to improve read quality and retained a final data of 398659457 read pairs, providing about 95.7% coverage.

### Comparative mapping with members of Family Canidae

The final selected reads of the dhole genome were compared with available genomes of domestic dog (*Canis familiaris*), African wild dog (*Lycaon pictus*) and dingo (*Canis lupus dingo*). Their genome information was downloaded from NCBI (Table 1). The reference genomes were indexed using Burrows-Wheeler Aligner (BWA) v 0.7.17 (Li et al. 2010) and the raw reads of dhole genome were aligned against the reference genomes using default parameters of BWA MEM module. The alignments from BWA MEM were streamed into SAMBLASTER v0.1.24 (Faust and Hall 2014) to exclude duplicates, add mated read tags and to separate out discordant, split read alignments using default parameters. The discordant and split read alignment sam files were converted into bam format using samtools 1.7. The mated alignments were then passed through Sambamba v0.6.6 (Tarasov et al. 2015) in order to sort and merge the alignments from the two libraries.

**Table 1:** Statistics of the reference genomes of other species used for comparative mapping of Asiatic Dhole Genome

| Species | Accession no. | GenBank PMID | No. of Scaffolds | Genome length (Mb) |
| --- | --- | --- | --- | --- |
| Dingo<br>(*Canis lupus dingo*) | GCF_003254725.1 | QKWQ00000000.1 | 2,444 | 2,439 |
| Domestic dog<br>(*Canis familiaris*) | GCF_000002285.3 | AAEX00000000.3 | 3,310 | 2,410 |
| African wild dog<br>(*Lycaon pictus*) | GCA_001887905.1_LycPicSAfr1.0 | LPRB00000000.1 | 803 | 2,358 |

### Identification of SNPs and InDels

FreeBayes v1.1.0-46 (Garrison & Marth 2012) with parallel implementation was used to identify the SNPs and indels. We used stringent base and mapping quality filters as per FreeBayes standard option, where -m option sets the mapping quality and -q sets the Phred scaled base quality to exclude alleles if the supporting base quality is less than 10. Minimum haplotype length of 5 bases was taken to allow continuous matches and to improve the variant calling process. The variants were emitted in a VCF compliant format and then filtered using bcftools filter module v1.8 based on a minimal read depth of 10, minimum quality of 30 and a maximal read depth as recommended by Li (2014).

The variant annotations were done using snpEff v4.3 in case of variants identified using domestic dog and dingo as reference genomes. The variants called against the African wild dog were not annotated, since there is no available genome annotation.

### Identification of Structural and Copy Number Variants

To determine the structural and copy number variants we used the combined approach including both read pair and read count algorithm (Tattini et al. 2015). The variants were mined using lumpyexpress in LUMPY v 0.2.13 (Layer et al. 2014) to identify intra-chromosomal translocations (BND), inversions (INV), deletions (DEL) and insertions (INS) with probability curve. To perform the analysis we used the concordant sorted alignments, split and discordant bam files derived from mapping to the reference genome. The filtering was done depending on the variant depth. Structural variants were only considered if the locus was supported by a minimum depth of 12x reads using bcftools filter option

The copy number variants were obtained using CNVnator’s with docker image ‘mustxyk/ubuntu-cnvnator’ from 0.3.2 version of the tool (Abyzov et al. 2011). We used the “-unique” option in order to obtain “q0” score of the calls as per author’s recommendation. The entire process used a window size of 1000 bases to identify the variants, which were later filtered by taking the “q0” scores between 0-0.5. The annotations for the filtered variants were done separately by the R package intanSV.

### Identification of Simple Sequence Repeats

Consensus sequence based on the Domestic Dog genome was created using bcftools consensus by putting back the filtered SNVs and InDels. The consensus genome was further searched for mono, di, tri, tetra, penta and hexa nucleotide repeats using PERF v0.2.5 (Avvaru et al. 2017) and MISA. The minimum length of the repeats was fixed at 12 bases as recommended by Subramanian et al. (2003). The repeat search conditions are provided in Table 2. The repeat motifs were intersected using bedtools intersect with a reciprocal match of 75% to call the SSRs concordant amongst the two tools.

**Table 2:**
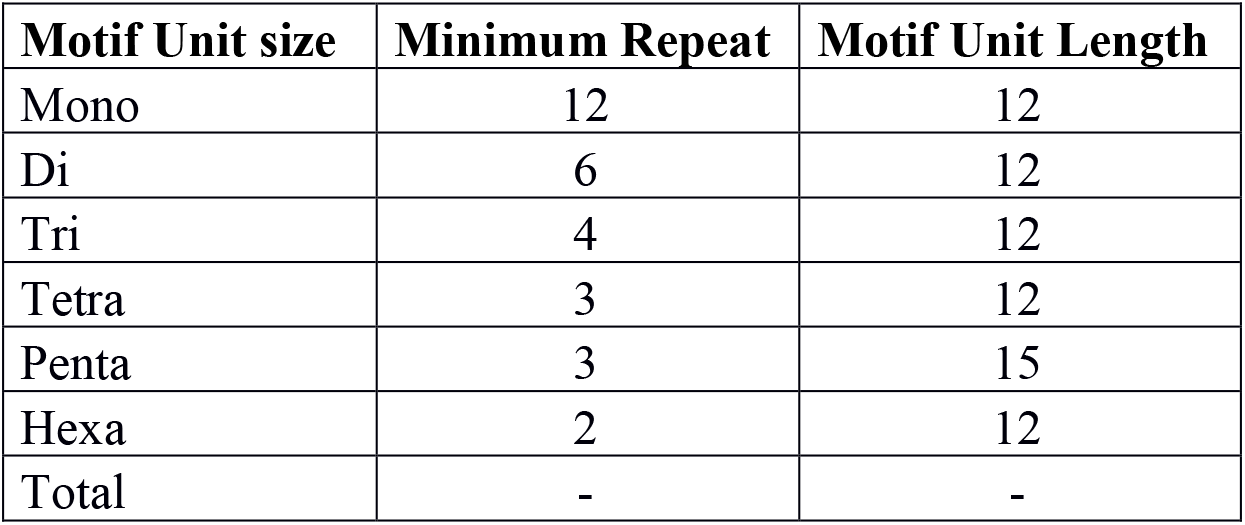
Motif unit and repeat length cut-offs

### Comparative mapping with dingo and dog for specific genes

To understand the peculiar differences in the dentition, coat colour and mammary glands in dholes we identified and compared the phenotypic genes using human and mouse as models with domestic dog and dingo. We transferred the phenotype to gene mapping from mouse and human to domestic dog and dingo based on orthologous relationship.

We obtained the existing orthology information between model organisms (human and mouse) and canines (dog and dingo) from ENSEMBL Biomart (Zerbino et al. 2018) and OMA orthology database (Altenhoff et al. 2018). The orthology information between model organisms and dhole were identified using bi-directional best PLAST (Parallel Local Sequence Alignment Search Tool) hit based on PLAST search (Nguyen et al. 2009). The protein sequences of mouse, human, dog, and dhole for PLAST search were obtained from ENSEMBL 2018 (Zerbino et al. 2018) and reference sequence database (RefSeq) at NCBI (O’leary et al. 2016) respectively.

Phenotype to gene mapping in human and mouse were obtained from mouse (Bello et al. 2015) and human phenotype ontology (Robinson et al. 2008), respectively. The gene ontology information for domestic dog was obtained from dog gene ontology ENSEMBL (Zerbino et al. 2018). The phenotypes to gene mapping related to coat colour, dentition and mammary gland function were transferred from reference model organisms (human and mouse) to domestic dog and dingo based on orthology relationship. We also mapped the gene ontology functions related to mammary gland function and development from domestic dog to dingo based on orthology relationship. In addition to that the known genes responsible for coat colour and dentition patterns in canines were obtained from the literature (Campana et al. 2016; Jernvall et al. 2012).

### Mitochondrial Genome Assembly

The reads mapping to the mitochondrial sequences from dingo, domestic dog and African wild dog genomes were extracted using samtools bam2fastq command. These reads were then error-corrected and assembled using SPAdes v3.11.1 (Bankevich et al. 2012) using the auto-multi-kmer mode.

## Results

### Variant calling across the reference genomes

From the 124.8 Gb data generated during dhole sequencing, we retained 398659457 reads after trimming low quality bases, adapters and discarding low quality sequences. In this data 99.16% of the clean reads were successfully mapped to the three reference genomes. We mined SNV, indels, structural and copy-number variant data and found ~13553269 SNV’s, ~2858184 indels, ~41000 SVs and finally about 1000 CNV in genomes of dhole, dingo and dog. Detailed statistics are tabulated in Figure 1 for all types of variants. From the many genes hosting the variants, we primarily looked at genes involved in dentition, pelage and mammary glands.

**Figure 1:**
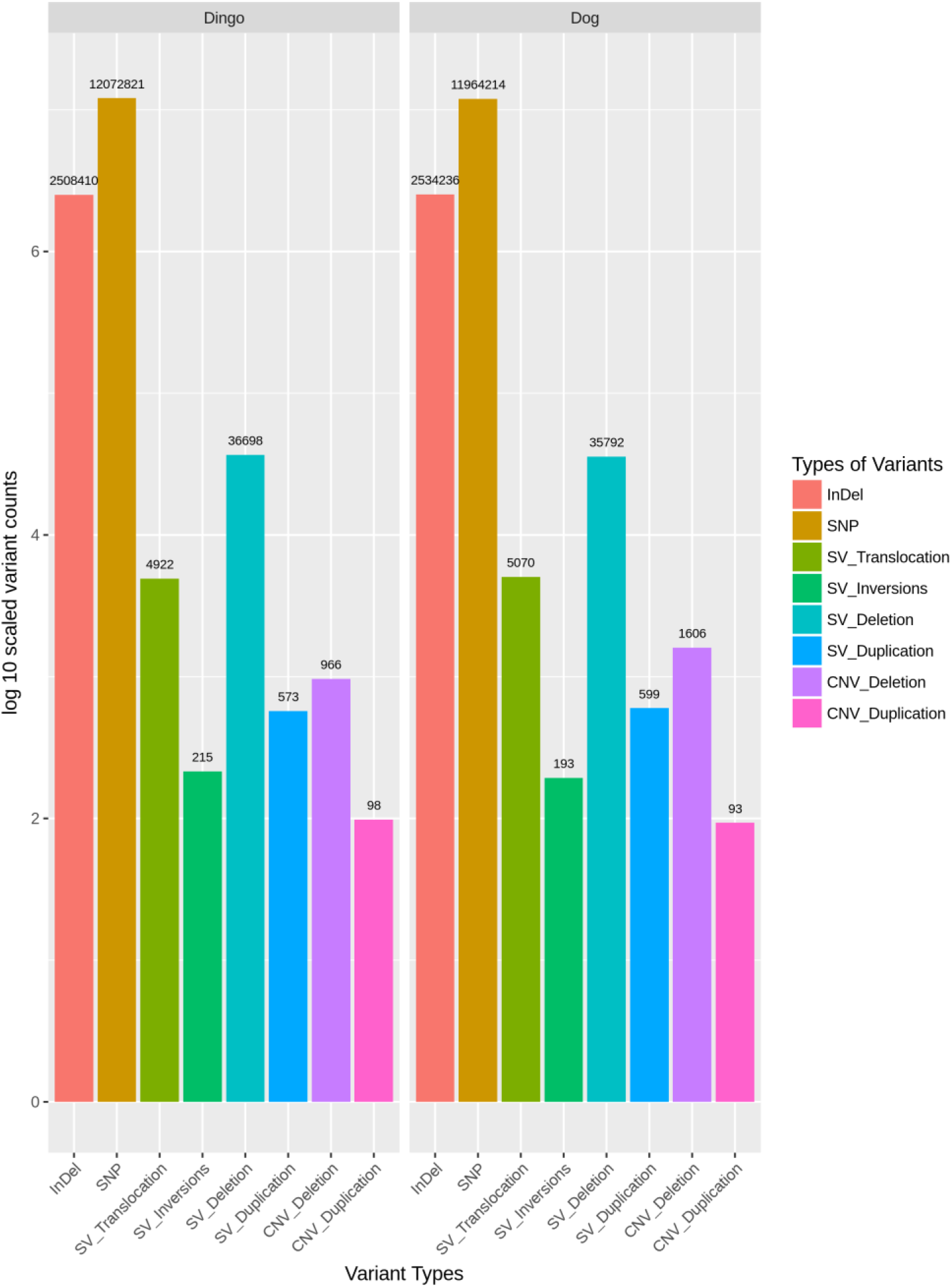
Frequency of structural variants in dhole using reference genomes of dingo and domestic dogs.

### SSRs in dhole based on domestic dog

Based on high quality SNPs and indels derived from filtering the FreeBayes emitted VCF, we created a consensus genome. We observed that 3.04% of the genomic length were part of SSRs in case of PERF whereas 3.65% was reported in case of MISA, with most of the contribution arising from hexa and tetra nucleotide SSRs (based on ∑(Length SSRs)*100/genome length). We observed a concordant 1854109 SSRs from 4692222 PERF predicted SSRs and 2813199 MISA predicted SSRs (Figure 2 & 3).

**Figure 2:**
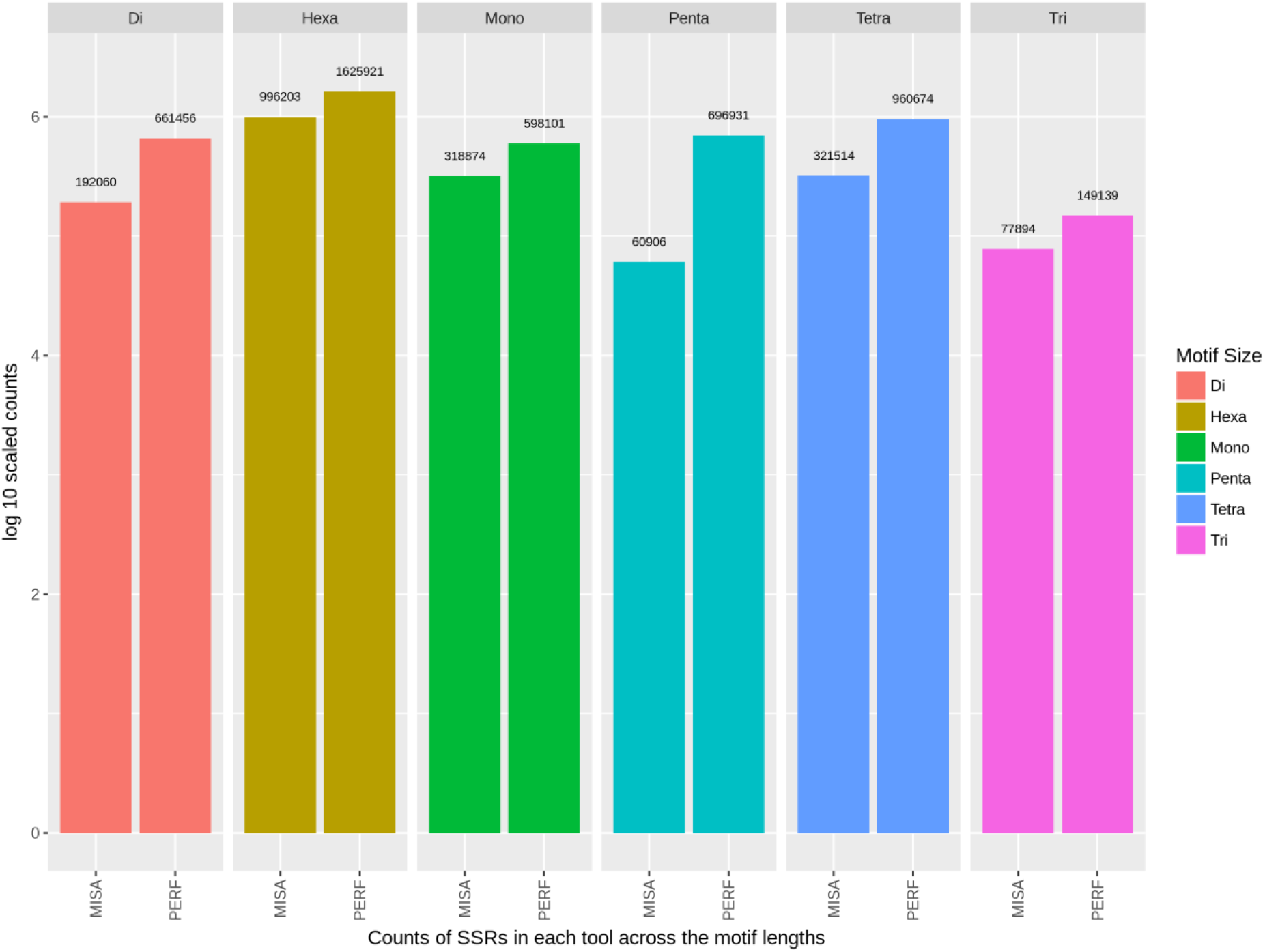
Distribution of SSR frequencies predicted using PERF

**Figure 3:**
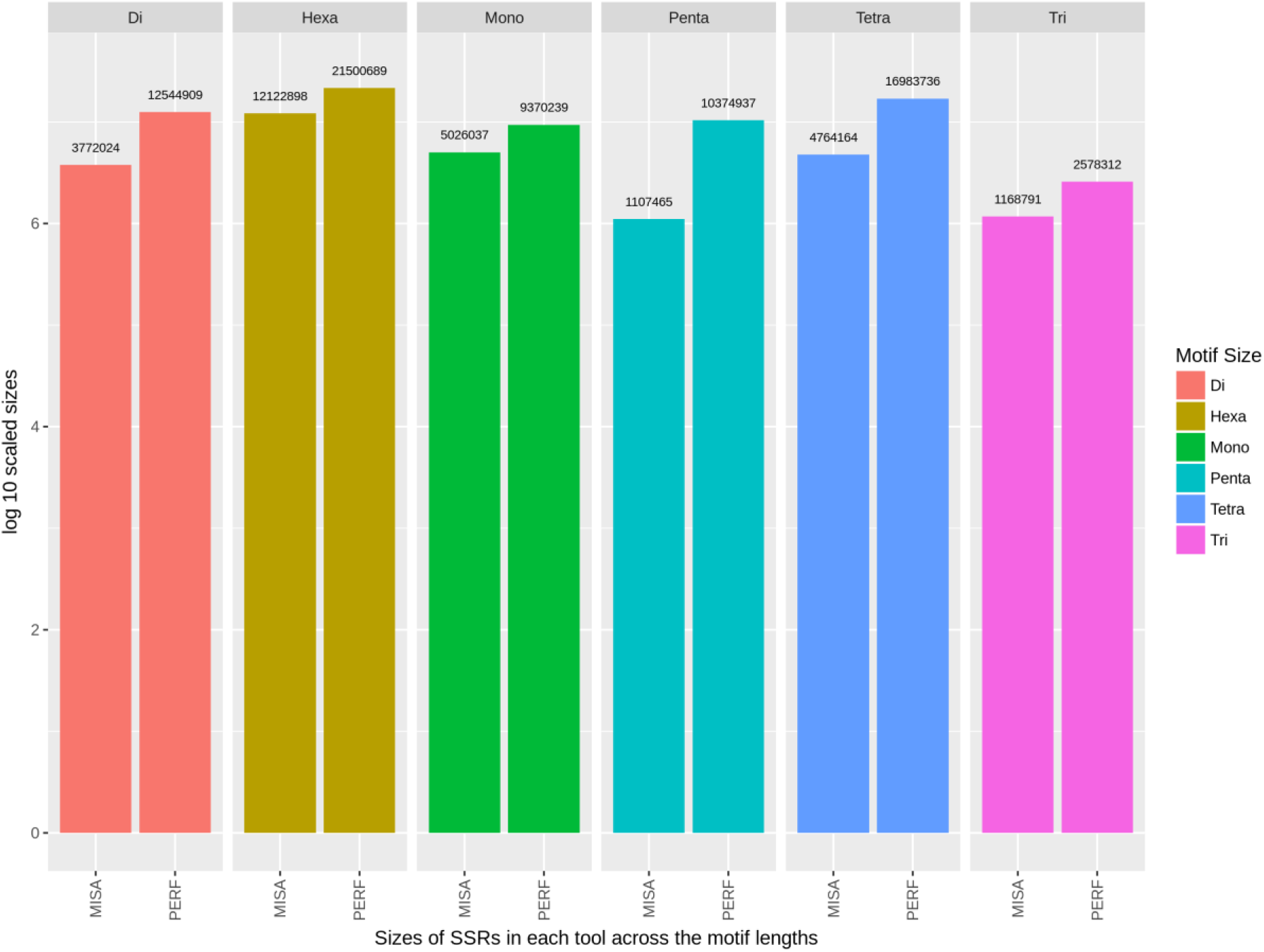
Distribution of SSR frequencies predicted using MISA

### Mitochondrial Genome Assembly

The baited sequences originating from the mitochondrial genomes of the canids were used to assemble the mitochondrial genome of Dhole into 37 scaffolds using SPAdes using the native multi-kmer approach. 36 scaffolds from the assembly had a sum length of 19,485 bases with a mean depth of 1.32x with the smallest scaffolds being 280 bases long. The GFA format pf the assembly was visualized using Bandage v0_8_1 (Wick et al. 2015), corroborated the same with the longest scaffold being circular having a depth of ~105.3x with a length of 16845 bases. The other 36 sequences were discarded and the circularized scaffold was annotated using Prokka v1.14-dev (Seemann 2014) using the Mitochondrial mode complying with GenBank recommendations. Annotation of the longest scaffold resulted in the prediction of 35 genes, 11 CDS and 24 tRNA.

### Variations in pelage, dentition and mammary gland related genes

We looked at the SNP variants in upstream region of genes coding for coat pattern, dentition and mammary glands. We selected the positively selected genes having higher ratio of non-synonymous vs synonymous mutations for further analysis. The top 50 upstream gene variants having SNPs were arranged in decreasing order for dhole vs. dingo and dhole vs. dog. The top ranked genes for coat pattern, dentition and mammary glands were found to play a role in signalling and developmental pathways. It is also important to compare the positively selected genes for the three traits with African wild dog being the first diverged genus before Cuon which will be a future work. (Figure 4,5,6).

**Figure 4:**
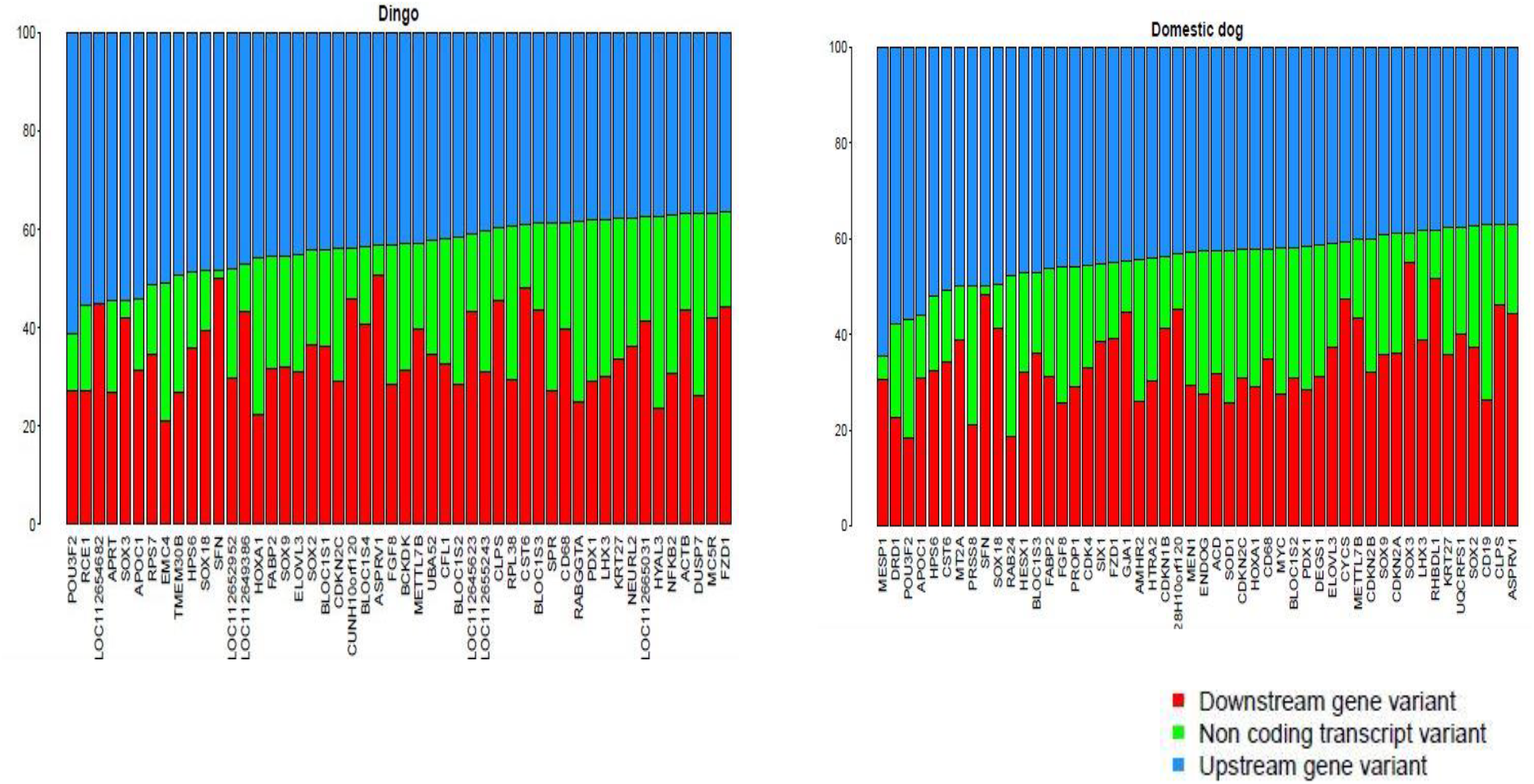
Percentage of variants distributed in upstream, non-coding and downstream regions of coat pattern genes

**Figure 5:**
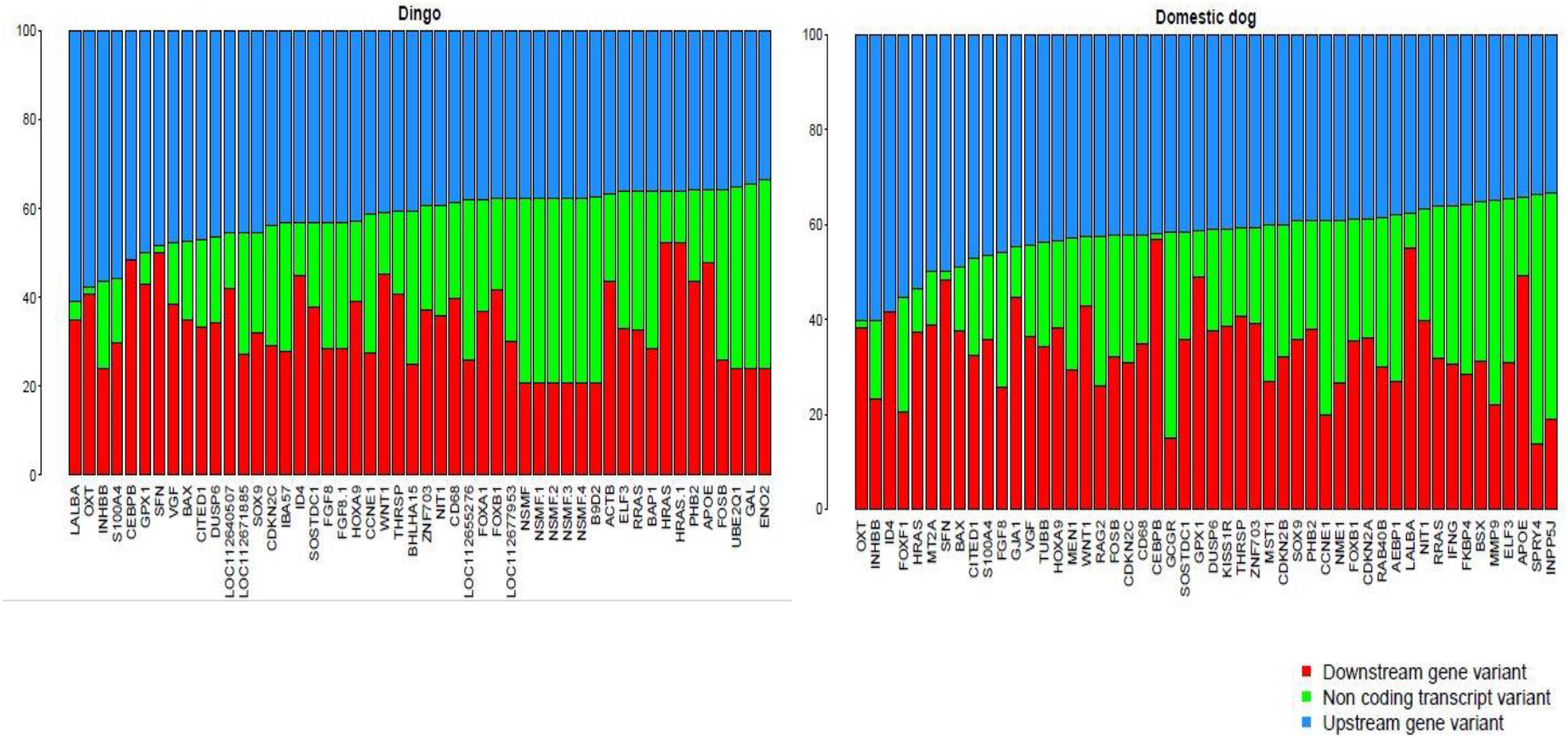
Percentage of variants distributed in upstream, non-coding and downstream regions mammary gland genes

**Figure 6:**
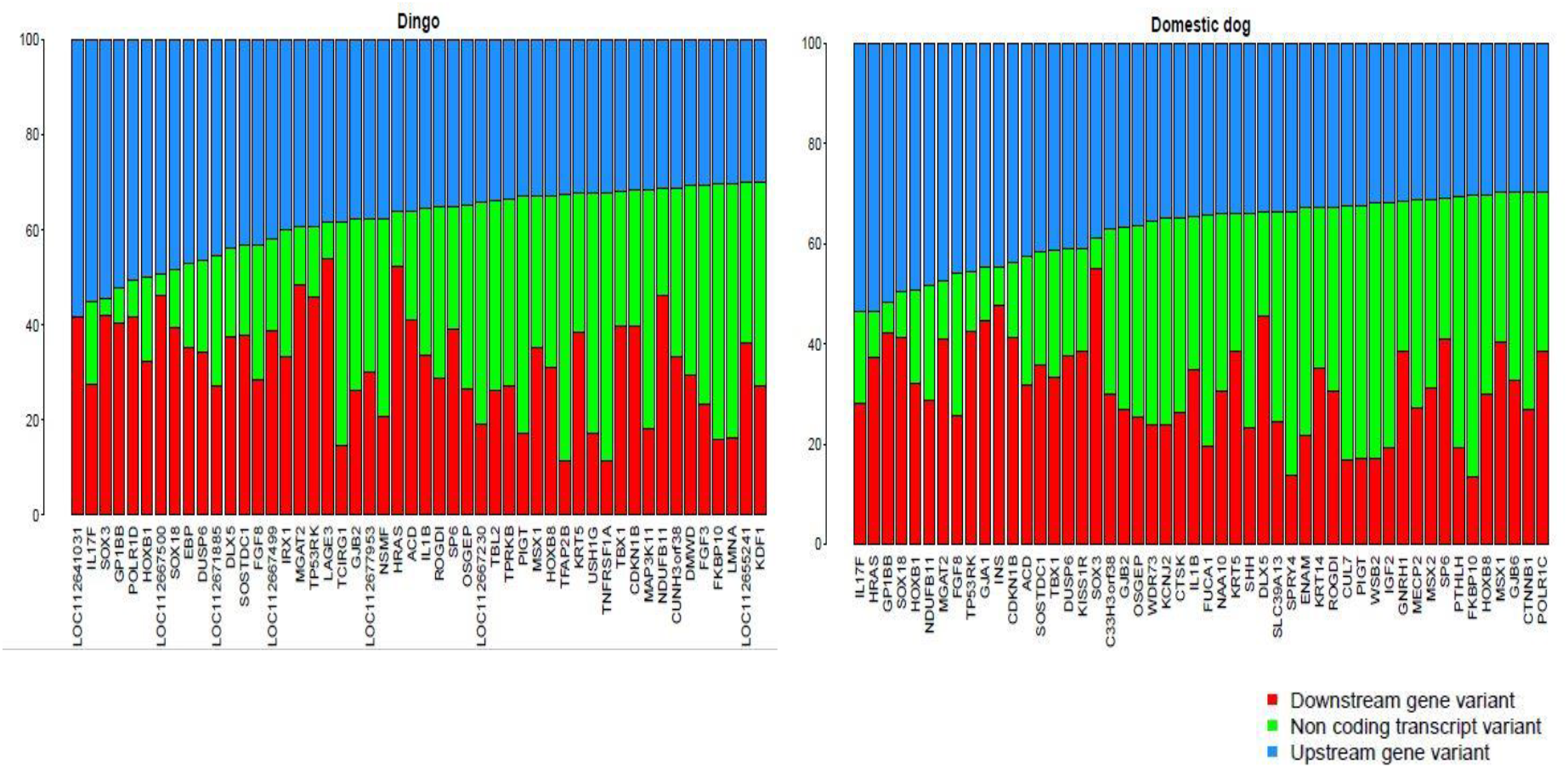
Percentage of variants distributed in upstream, non-coding and downstream regions dentition genes

We also considered nine different genes involved in pathways for melanin formation (ASIP, MC1R, MLPH, MITF, PMEL, TYRP1) and hair growth and patterns (FGF5, KRT71, RSPO2) (Campana et al. 2016) to understand the variations in dholes coat pattern and their expression in comparison to the Dingo and domestic Dog. The gene DEFB103A is absent both Dingo and domestic Dog. We found that genes like MITF (responsible for white spotting phenotypes in dog and coat colour variants), PMEL (responsible for merle pattern) and ASIP (responsible for darker and lighter hair colours) are not showing any evident variation in exon regions and showing most of the variation in upstream promoter region and downstream regions and non-coding regions.

While in case of MC1R gene, it is highly polymorphic in upstream promoter and downstream regions. The polymorphisms could reduce the ability of the melanocortin 1 receptor to stimulate eumelanin production, causing melanocytes to make mostly pheomelanin which supports the fact that Asiatic wild dog has red coat colour. (Figure 7)

**Figure 7:**
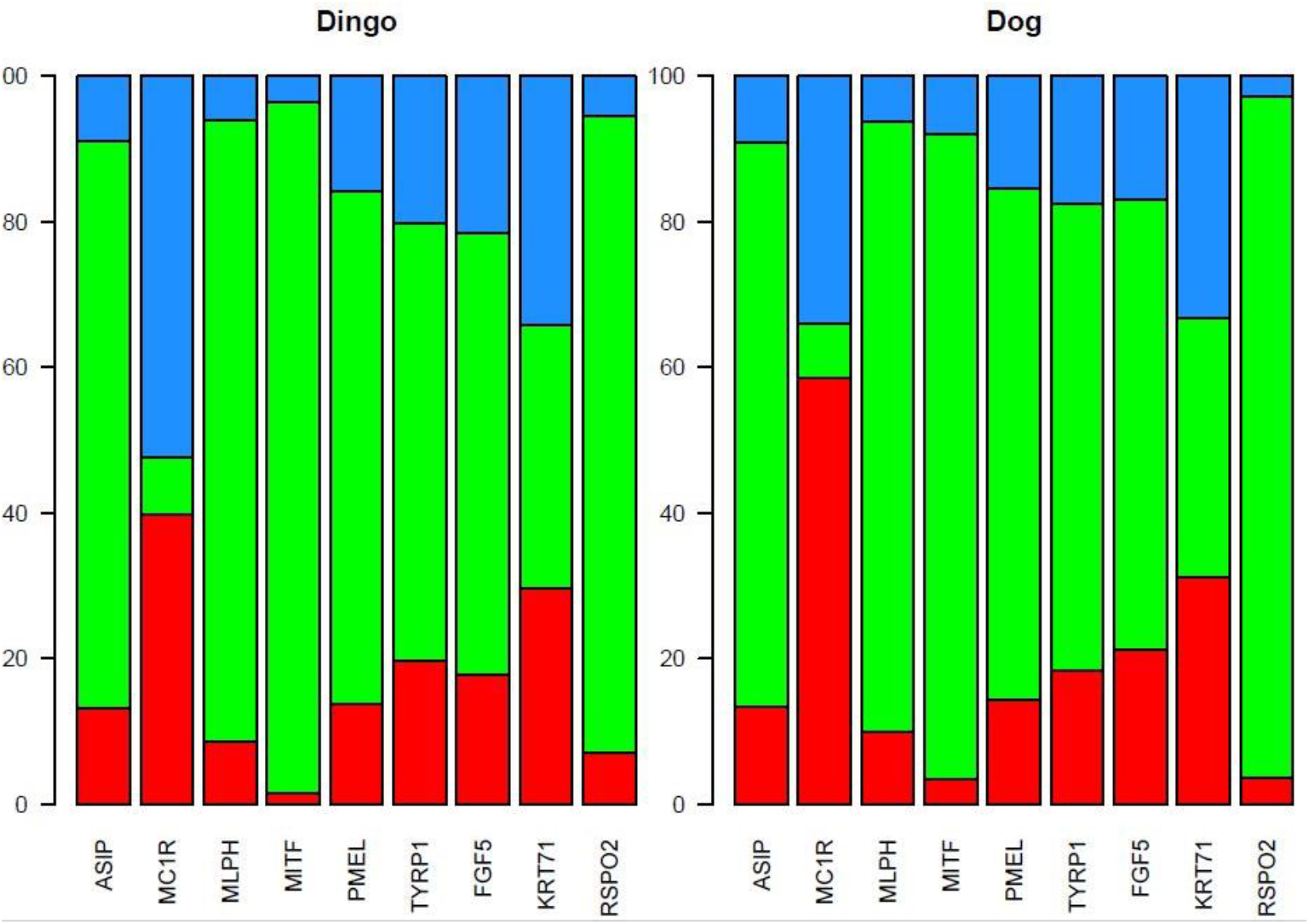
Percentage of variants distributed in upstream, non-coding and downstream regions known genes of coat pattern

## Conclusion

This is the first reported genomic study of the elusive, social Asiatic wild dog with an exploratory comparison with related canids dingo, African wild dog and domestic dog. This research yielded a draft genome survey of 2.4 Gb with 52X coverage basing on the domestic dog genome. This work is mainly focused on identification of structural variants, copy number variants, simple sequence repeats and single nucleotide polymorphisms and its comparison with other canid species to develop an evolutionary insight for this monophyletic genus. This will also help in understanding the divergence of two monophlyletic genomes *Cuon* and *Lycaon* during the course of evolution and differences arose in Dhole as compared to other canids in the form of coat pattern, dentition and mammary glands.

## Acknowledgement

This research is a collaborative work between Wildlife Institute of India and Nucleome Informatics Private Limited. The authors acknowledge the support from Maharashtra State Forest Department for the permits to conduct this work. We thank the forest department officials, staff and our field assistants Roshan and Akshay for their assistance in fieldwork during dhole capture and blood sampling. Our sincere thanks to the Dean, Faculty of Wildlife Sciences, Director and Research Co-ordinator of WII for their support. This research was funded by Maharashtra State Forest Department and National Tiger Conservation Authority, Government of India.

